# Hippocampal recruitment of cortical engrams underlies remote memory generalization

**DOI:** 10.64898/2026.04.30.721635

**Authors:** Gabriel Berdugo-Vega, Irene Ferreres, Jules Orsat, Myriam Schioppa, Johannes Gräff

## Abstract

Time-dependent generalization of remote fear memories is generally viewed as a passive and progressive loss of contextual precision, yet its underlying mechanisms remain poorly understood. Here, we investigated the neurobiological correlates of remote generalization using a combination of engram and projection-specific manipulation technologies in mice. Our results show that remote fear generalization results in an increased recruitment of learning-associated engrams into recall ensembles in the medial prefrontal cortex (mPFC), a crucial brain area for remote memory storage. Moreover, we find that this recruitment, and concomitantly remote generalization, requires ventral hippocampal (vCA1) inputs conveying contextual information to the mPFC, for which activity of parvalbumin-expressing interneurons proves necessary. Together, our findings suggest that time-dependent fear generalization arises from active hippocampal-prefrontal circuit mechanisms rather than passive loss of contextual information.

## INTRODUCTION

Time-dependent generalization is a core hallmark of remote memories. In the context of fear, generalization refers to the process by which an innocuous stimulus or context is perceived as threatening because of similarities with a previous experience. While generalization can promote survival and aid flexible behaviour ^1^, for instance by coordinating fear responses to stimuli similar to the original threat experience, overgeneralization can also be maladaptive. This is the case in anxiety and post-traumatic stress disorders where exacerbated fear responses are triggered in non-threatening conditions ^2^. Time-dependent generalization is generally regarded as a progressive loss of memory precision ^3^, but its neurobiological mechanisms are not well understood.

Episodic fear memories are first processed in the hippocampus and later transferred and consolidated into cortical circuits for long-term storage ^4^, during which contextual detail is believed to be lost resulting in a gist-like nature of cortically-dependent remote memories and generalization ^5,6^. Consequently, neural correlates of time-dependent generalization have been studied, among others, in hippocampal and cortical areas ^7^. Interestingly, two studies using lidocaine or chemogenetic inhibition of the ventral portion of CA1 (vCA1) revealed decreased remote fear expression in a novel context ^8,9^, suggesting that vCA1 might be important for remote memory generalization. vCA1 has divergent projections to key structures involved in cognition ^10^, including to the mPFC ^11,12^, a crucial structure for the storage and retrieval of remote memories that has previously been linked to generalization as well ^8,9,13^. The vCA1–mPFC projection is involved in anxiety ^14–16^, social ^17,18^ and spatial memory ^19^, as well as fear extinction ^20,21^, but its role for remote memories has not been investigated.

In spite of significant advances in memory research with the discovery and manipulation of engrams (i.e., cells active during learning and responsible for recall) including in the mPFC that control remote memory recall ^22–25^, few engram studies have focused on time-dependent memory generalization ^26,27^. How the proposed loss in remote memory precision manifests at the level of fear engrams in the cortex and the potential involvement of the hippocampus during generalization remain thus unknown. In this study, we investigated these questions from an engram perspective and with a particular focus on the vCA1–mPFC projection.

## RESULTS

### Remote generalization is associated with recruitment of mPFC engrams and vCA1 inputs

We first aimed at finding cortical engram correlates of time-dependent memory generalization. For this, we used TRAP2-Ai14 mice ^23^, in which tamoxifen administration after fear conditioning results in the recombination of a STOP cassette in the ROSA26 locus and thereby drives the expression of tdTomato (Tom) specifically in the cells active during encoding (Fig. 1A). This allows for the permanent tagging of fear memory-involved cells (i.e., A engrams). Two days or two weeks later (recent or remote time, respectively), mice were re-exposed to the same (AA) or a different (AC) context and freezing quantified. Animals were perfused 90 minutes after recall for quantification of cFos immune-positive cells, which represents the memory recall ensemble. We observed that, while mice tested in the same context displayed similar levels of freezing at recent and remote recalls, mice tested in the novel context C showed increase in freezing specifically at remote times, indicative of time-dependent memory generalization (Fig. 1B).

**Figure 1:**
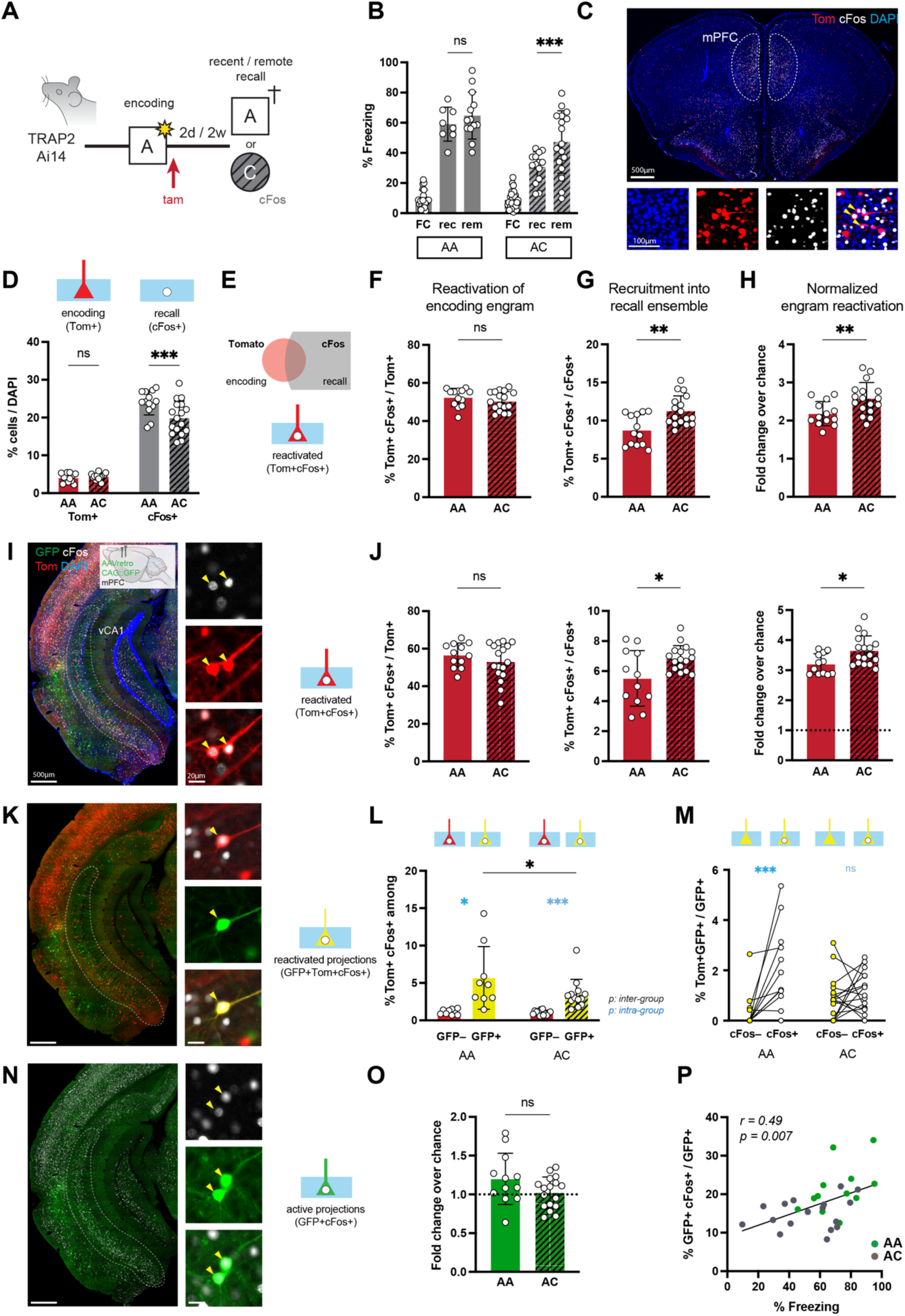
Remote generalization is associated with recruitment of mPFC engrams and vCA1 inputs. **A.** Scheme depicting the experimental design for the study of time-dependent remote memory generalization in TRAP2-Ai14 mice, consisting of a contextual fear conditioning paradigm followed by either recent (2 days) or remote (2 weeks) recall in the same -A- or a novel context -C-. This transgenic mouse line allows for the tagging of fear memory engrams upon tamoxifen (tam) administration at the time of conditioning. **B.** Percentage of freezing at recent and remote recall of animals exposed to the same (AA) or to the novel (AC) context, showing time-dependent increased freezing in the novel context indicative of generalization. **C.** Immunofluorescence picture with magnifications of learning (tdTomato, Tom^+^), recall (cFos^+^) and reactivated (Tom^+^cFos^+^, yellow arrowheads) ensembles in the mPFC of animals sacrificed after remote memory recall. **D – H.** Cellular quantifications in the mPFC including cell densities (D) and engram reactivation (as depicted in E) of AA and AC mice. Reactivation is showed as percentage of the encoding engram (Tom^+^cFos^+^/Tom^+^, F), recruitment into the recall ensemble (Tom^+^cFos^+^/ cFos^+^, G) and total normalized (fold change over chance levels, H). **I – J.** Immunofluorescence and magnifications (I) for the assessment of engram reactivation (yellow arrowheads) and quantifications (J) in vCA1 of AA and AC mice, as in E – H. Note the increase in recruitment and total fear engram reactivation in both the mPFC and vCA1 of AC mice. Here, vCA1 neurons projecting to the mPFC can be observed upon injection of a GFP-retrograde tracer in the mPFC, as shown in I. **K – M.** Immunofluorescence and magnifications (K) for the assessment of reactivated projections (GFP^+^Tom^+^cFos^+^, yellow arrowheads), with quantifications of engram reactivation among projecting or non-projecting cells (GFP^+^ or GFP^−^, respectively; L) and of cFos expression among originally traced engrams (Tom^+^GFP^+^; M) in vCA1 of AA and AC mice. **N – O.** Immunofluorescence and magnifications (N) for the assessment of active projections (GFP^+^cFos^+^, yellow arrowheads) and quantifications (O), showing no differences in total activity of vCA1–mPFC projections between AA and AC mice. **P.** Correlation between freezing and the percentage of active projections across AA (green) and AC (grey) mice. Data are represented as bar graphs (filled or stripped for AA and AC mice, respectively) with individual animal values (dots; N > 7 for behaviour and cellular analysis) and as correlation (P). Error bars indicate SD. Black and blue significance refers to inter- and intra-group comparisons, respectively. Scale bars = 500, 100 and 20 μm as indicated. *** *p* < 0.001; ** *p* < 0.01; * *p* < 0.05; ns, non-significant, by tests as indicated in Table S1.

In order to understand how generalization impacts engram cell reactivation in the mPFC, we quantified the numbers of Tom^+^, cFos^+^ and double positive cells in this area of mice tested at remote times (Fig. 1C). While there were no differences in the number of encoding (Tom^+^) cells, we observed a significant decrease in the number of cFos^+^ neurons in animals exposed to the C context (Fig. 1D), indicative of a smaller recall ensemble. To account for differences in ensemble size, reactivated cells can be normalized to the encoding, recall or both (vs. chance levels) ensembles (Fig. 1E). Reactivation of the original encoding engram (Tom^+^cFos^+^ / Tom^+^) remained unchanged between groups suggesting that the same percentage of engrams are reactivated after recall in either context A or C (Fig. 1F). However, recruitment of encoding engrams into the recall ensemble (Tom^+^cFos^+^ / cFos^+^) was increased in the AC group, indicating a greater contribution of reactivated fear cells to the representation of context C (Fig. 1G). This was explained by the overall reduction in activity observed in non-engram cells of AC mice (Fig. S1A). Reactivation was significantly above chance levels in both groups, and the fold change over chance was increased in AC mice (Fig. 1H), supporting the notion of a higher contribution of fear engrams to the cortical representation of context C. These results suggest that generalization is associated with changes in engram cortical activity that result in a higher recruitment of fear engrams when representing a novel context C.

Memory generalization has been previously linked to vCA1 function ^8,9^, but a potential role of the vCA1–mPFC projection in remote memory has never been studied. This prompted us to use a GFP-retrotracer injected into the mPFC of AA and AC mice (Fig. 1I) in order to investigate activity patterns of the vCA1–mPFC projection during generalization. A brain-wide assessment of retrogradely traced GFP somas revealed canonical prefrontal inputs, such as the basolateral amygdala, retrosplenial and entorhinal cortices, as well as vCA1 (Fig. S1B). Cellular quantifications showed no differences in the number of vCA1 traced or recombined cells between AA and AC animals, although again a reduction in the number of cFos^+^ cells in AC mice was observed (Fig. S1C). Next, we performed an analysis of whole-vCA1 engram reactivation in AA vs. AC animals similar to that in the mPFC. Interestingly, we found that the three reactivation parameters changed in the same magnitude and direction as they did in the mPFC (Fig. 1J), with strong and significant correlations between vCA1 and mPFC (Fig. S1D), suggesting that engram activity between these two areas is synchronized. Importantly, this was not the case for other hippocampal structures such as the DG, where reactivation decreased to chance levels at remote times (Fig. S1E), as also shown by others ^22,27^. Subsequently, we studied engram reactivation within the vCA1–mPFC projection itself (Fig. 1K). This revealed that the proportion of reactivated cells was greater among GFP^+^ vs. GFP^−^ cells for both AA and AC mice (Fig. 1L), suggesting that engrams are enriched in the vCA1–mPFC projection. More specifically, within the projection, reactivation in AA was greater than in AC mice (Fig. 1L), indicating contextual specificity that was not observed when quantifying reactivation in the entire vCA1 area as a whole (Fig. 1J vs. 1L). Supporting this, we found that from all originally traced encoding fear engrams (Tom^+^GFP^+^) those of AA animals were more likely to reactivate than those of AC animals (Fig. 1M). The remaining cells of the vCA1-mPFC projection (Fig. 1N) showed no differences in activity (GFP^+^cFos^+^, total vs. chance) among groups (Fig. 1O).

Finally, we investigated whether histological data alone was sufficient to separate groups in a dimensionality reduction approach. For this, we performed linear discriminant analysis from hippocampal and cortical cellular parameters of animals tested at remote times and, additionally, a small sample of animals tested and quantified for the same parameters after recent recall. We observed that AA and AC recent recall mice were the only clearly separated groups whereas AA and AC remote recall mice overlapped among themselves and with recent recall groups (Fig. S1F). When focusing on the AC remote group, we observed that 4 out of the 6 mice with the lowest freezing values overlapped with the AC recent group, whereas mice with higher freezing values predominantly overlapped with the AA remote group, suggesting that AC remote samples segregate based on the presence of generalization. This is remarkable considering that freezing was not used as input variable during the analysis. Interestingly, engram reactivation parameters preferentially influenced separation between recent and remote AA groups, whereas the activation of the vCA1-mPFC projection was the variable most strongly influencing separation within recent and remote AC mice. Supporting this, freezing response was significantly correlated with engram parameters at recent, and with projection parameters at remote times (Fig. S1G, top row). This suggests that the relative importance of engrams vs hippocampal-cortical projections might shift as a function of consolidation. Moreover, from all cellular parameters analysed, the only one significantly correlated with freezing at remote recall was the proportion of active vCA1–mPFC projections (Fig. 1P and S1G, right), suggesting that hippocampal inputs might act as modulators of remote memory expression.

### vCA1–mPFC projections are necessary for remote generalization

To causally evaluate whether inputs from vCA1 to mPFC are necessary for remote memory generalization, we used a conditional chemogenetic approach based on the injection of a retrograde Cre virus in the mPFC and Cre-dependent inhibitory DREADD (hM4Di-mCherry)-containing virus in vCA1. This allows for the expression of hM4Di-mCherry exclusively in vCA1 cells projecting to the mPFC, which can be inhibited upon administration of the synthetic agonist CNO (Fig. 2A). Since this approach prevents the use of Cre-based TRAP2 mice to label fear engrams, *cFos::tTA* mice ^28^ and TRE::GFP viruses were used instead, and doxycycline (DOX) removed two days before memory encoding. After fear conditioning in A, DOX was immediately re-introduced in drinking water to close the engram labelling window. Two weeks later, mice were administered with CNO or its vehicle (VEH) 30 minutes before memory recall in either context A or C. Similarly to TRAP2 mice, *cFos::tTA* mice administered with VEH showed contextual generalization and comparable freezing in A and C. However, we found that CNO administration reduced freezing by about 25% specifically in AC but not AA animals, indicating that vCA1–mPFC projections are necessary for remote memory generalisation (Fig. 2B).

**Figure 2:**
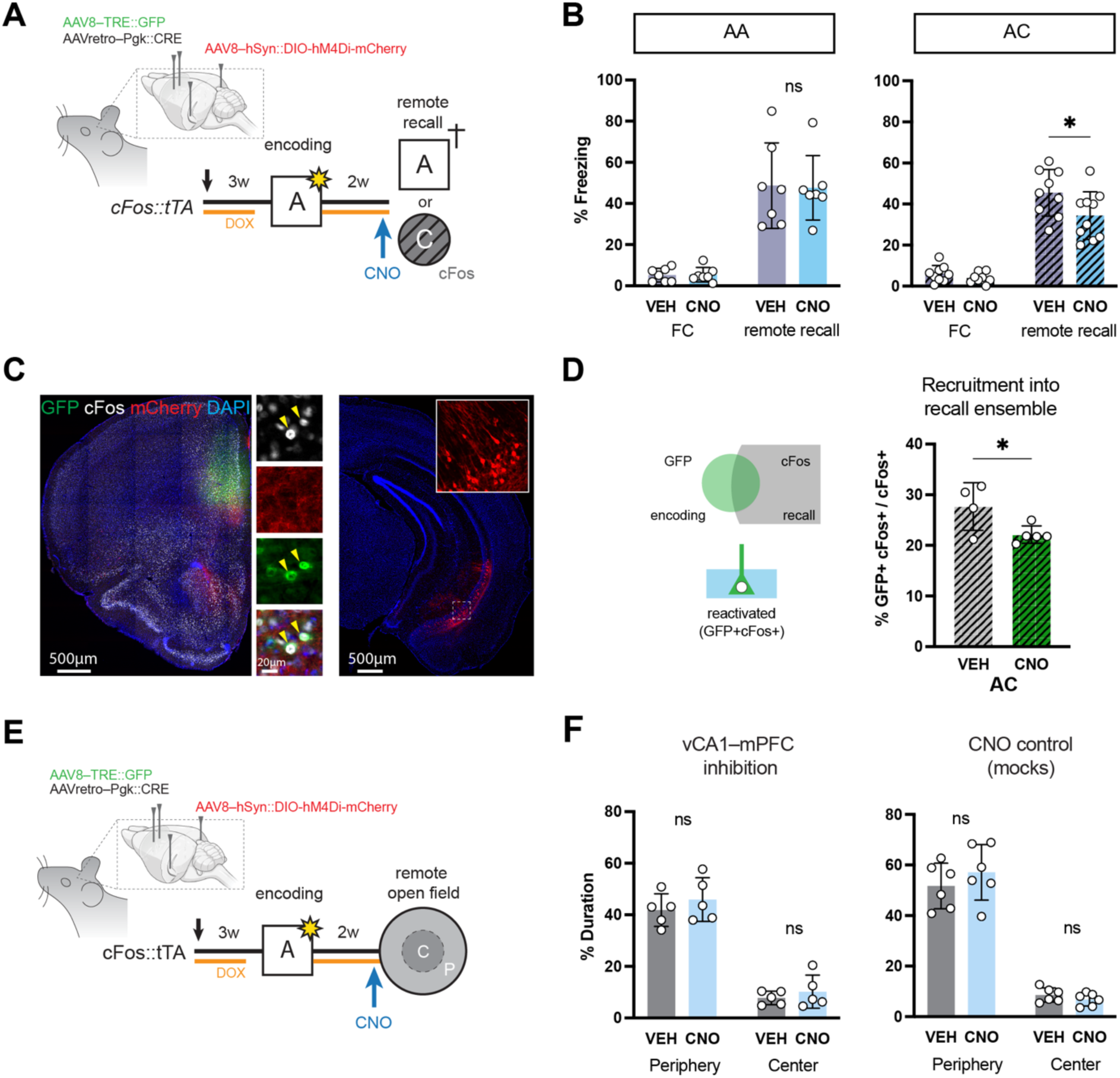
vCA1–mPFC projections are necessary for remote generalization. **A.** Scheme depicting the experimental design for the manipulation of the vCA1–mPFC projection during memory generalization. In this setup, the indicated combination of viruses allows the selective expression of hM4Di-mCherry in mPFC-projecting vCA1 neurons, which can then be inhibited upon CNO administration before remote recall in mice exposed to either the same conditioned (AA) or a novel (AC) context. The use of TRE::GFP in *cFos::tTA* mice additionally allows the doxycycline (DOX)-dependent labelling of mPFC fear engrams at the time of learning. **B.** Percentage of freezing in AA and AC mice injected with either a vehicle (VEH) or CNO, showing a reduction in memory generalization specifically in CNO-injected AC mice. **C – D.** Immunofluorescence picture with magnifications (C) of learning (GFP^+^), recall (cFos^+^) and reactivated (GFP^+^cFos^+^, yellow arrowheads) mPFC ensembles, together with fibers from vCA1–mPFC projecting neurons (mCherry^+^) and quantification of reactivation (D), showing decreased recruitment into recall ensembles in CNO as compared to VEH injected AC animals. **E – F.** Scheme depicting the experimental design for the study of stress-related behaviour 2 weeks after fear conditioning (E) and quantifications (F), showing lack of effect of vCA1–mPFC projection inhibition in *cFos::tTA* mice (F, left) as well as of CNO administration on non-injected animals (F, right) during the exploration of an open field. Data are represented as bar graphs (filled or stripped for AA and AC mice, respectively) with individual animal values (dots; N > 6 for fear memory, > 4 for stress behaviour and > 3 for cellular analysis). Error bars indicate SD. Scale bars = 500 and 20 μm as indicated. * *p* < 0.05; ns, non-significant, by tests as indicated in Table S1. See also Figure S2

Analysis of these brains revealed mCherry expression in vCA1 somas as well as puncta and fibers in the mPFC area, as expected (Fig. 2C). Quantifications in the mPFC showed no differences in the number of GFP^+^ or cFos^+^ cells upon CNO administration in AA and AC animals (Fig. S2A), and no effect of CNO in the reactivation of the original encoding engram (Fig. S2B, left). Importantly, however, CNO led to a significant reduction in the recruitment of the fear engram to the recall ensemble (GFP^+^cFos^+^ / cFos^+^) in AC mice (Fig. 2D), that was not observed in AA mice (Fig. S2B, right). These results indicate that hippocampal inputs regulate activity re-allocation in the mPFC, resulting in reduced overlap of fear and recall ensembles in a novel context that is associated with decreased generalization.

A number of controls confirmed these results. First, the absence of a behavioural phenotype in AA mice suggests that the effect of inhibiting the projection is context-specific, and not the result of a technical artifact derived from CNO administration. Second, regarding the effects at the cellular level in engram recruitment, we took advantage of sections from CNO-administered animals that were mis-injected and thus excluded from our previous analysis. Hemispheres where no mCherry signal could be detected in the mPFC (i.e., where the hemilateral projection was not inhibited) did not show a reduction in engram recruitment when matched to the contralateral, mCherry^+^ side, confirming that the effect in activity re-allocation is specific to the inhibition of the projection (Fig. S2C). Third, given the proposed role of vCA1–mPFC projections in regulating anxiety ^14–16^, we tested whether our reported reduction in freezing could be the result of reduced overall stress. For this, we inhibited the vCA1–mPFC projection prior to an open field test (Fig. 2E). Analysis of the time spent in the centre vs. the periphery did not show any differences between CNO and VEH administered groups, and neither did we find differences in mice of the same genotype that were not virally injected (i.e., to control for the effect of CNO alone) (Fig. 2F). Velocity and distance travelled were also unaffected (Fig. S2D). Therefore, these control experiments confirm that the vCA1–mPFC projection promotes remote contextual generalization through recruitment of cortical fear engrams into the recall ensemble.

### vCA1 routes contextual information of the novel context during remote generalization

Considering that inhibition of the vCA1–mPFC projection reduces freezing in the novel context C but not in A (Fig. 2), and that vCA1 routes context-specific information to the mPFC (Fig. 1L and 1M), we reasoned that the necessary hippocampal input to the cortex that drives generalization should be the contextual representation of the novel context C. To test this, we transitioned from a projection- to an engram-inhibition approach by injecting Cre-dependent hM4Di-mCherry in the vCA1 of TRAP2 mice. In this case, tamoxifen administration results in the conditional expression of hM4Di-mCherry in vCA1 engram cells for later inhibition via CNO-administration prior to the remote exposure to the context C. We tested independent cohorts of mice in a variation of our previous behavioural paradigm, in which an exposure to the context C precedes fear conditioning (5 days, to avoid memory linking) (Fig. 3A). Here, tamoxifen administration followed either exploration of the non-fear associated context C or, alternatively, fear conditioning in the context A. We also included a cohort of control mice in which tamoxifen administration occurred in the home cage the day before the first C exposure, to control for both CNO administration in the TRAP2 line as well as for the effect of the prior exposure to context C. Behavioural analyses on the day of conditioning revealed that the previous exposure to context C did not affect exploration of the context A (Fig. S3A). At remote recall, only inhibition of the vCA1 representation of context C but not of the context A or home cage reduced generalization when compared to each respective VEH group (Fig. 3B). Inhibition of “A” engrams did not prevent recall in conditioned context A either (Fig. S3B), suggesting that the hippocampal “A” representation is no longer needed for remote recall. These results confirmed that hippocampal input of the non-fear associated context, but not the original fear-associated one, is necessary for remote memory generalization.

**Figure 3:**
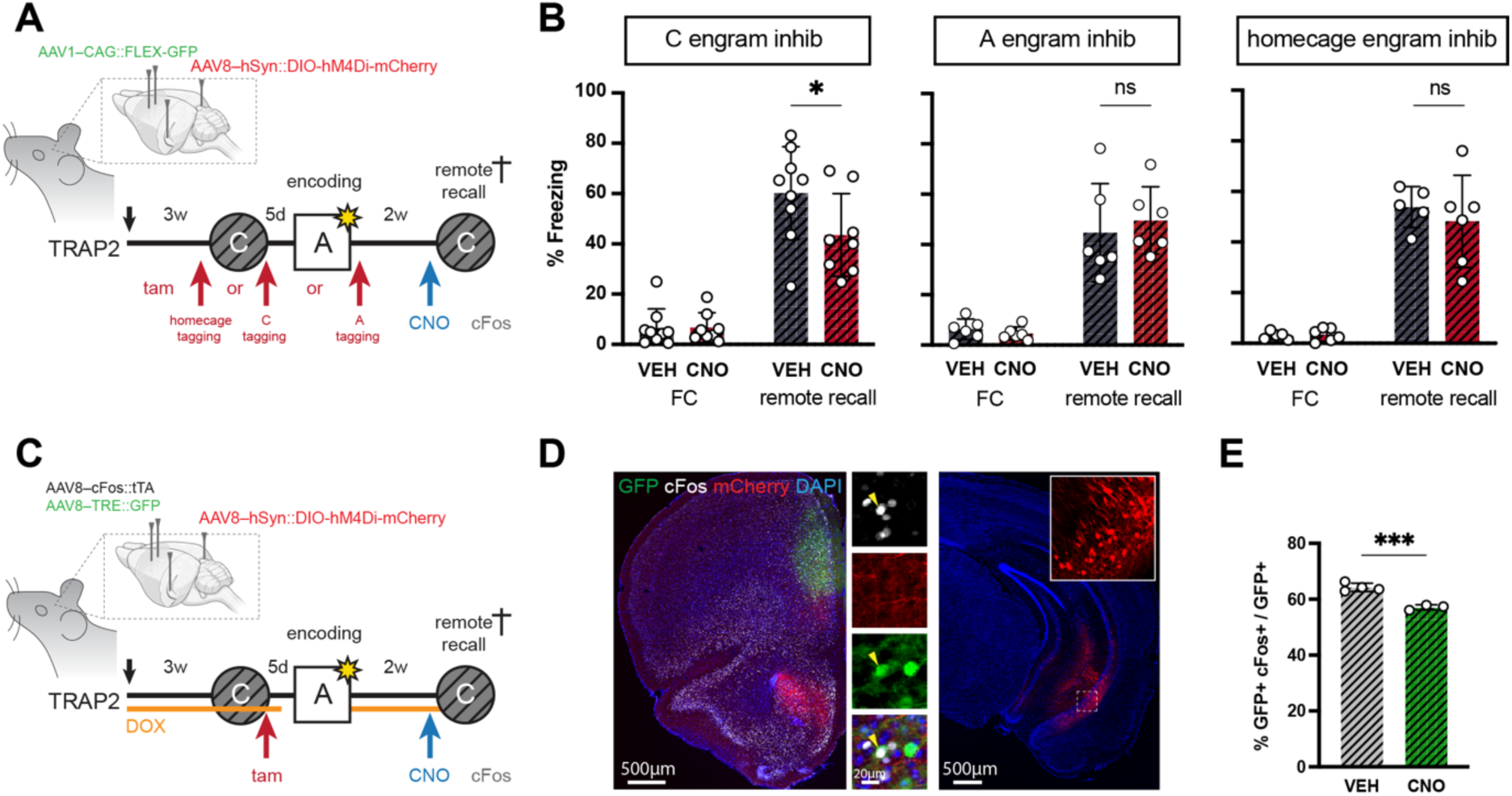
vCA1 routes contextual information during remote generalization. **A.** Scheme depicting the experimental design for the manipulation of vCA1 engrams during memory generalization. In this setup, the indicated combination of viruses in TRAP2 mice allows the selective expression of hM4Di-mCherry via tamoxifen-dependent recombination in vCA1 neurons. Tamoxifen was administered at different times in 3 different cohorts of mice, including after exploration of either the homecage, the context C or the conditioning context A for the trapping, and later inhibition via CNO administration before remote recall, of contextually-specific vCA1 neuronal ensembles. **B.** Percentage of freezing in AC mice injected with either a vehicle (VEH) or CNO, showing a reduction in memory generalization specifically after “C” (but not “A” or homecage), engram inhibition. **C.** Scheme depicting the experimental design for the manipulation of vCA1 engrams “C”, and additionally including the injection of cFos::tTA and TRE::GFP for the DOX-dependent labelling of context “A” mPFC engrams. **D – E.** Immunofluorescence picture with magnifications (D) of “A” (GFP^+^), recall (cFos^+^) and reactivated (GFP^+^cFos^+^, yellow arrowheads) mPFC ensembles, together with fibers from vCA1–mPFC projecting neurons (mCherry^+^) and quantification (E) showing decreased reactivation of fear “A” mPFC engrams upon inhibition of vCA1 “C” engrams. Data are represented as bar graphs with individual animal values (dots; N > 4 for fear memory, > 2 for cellular analysis). Error bars indicate SD. Scale bars = 500 and 20 μm as indicated. *** *p* < 0.001; * *p* < 0.05; ns, non-significant, by tests as indicated in Table S1.

To confirm a decreased contribution of cortical fear “A” engrams to generalization while at the same time being able to inhibit vCA1 “C” engrams, we further injected TRAP2 mice with cFos::tTA + TRE::GFP viruses for DOX-dependent labelling of mPFC engrams at the moment of conditioning in A (Fig. 3C). Despite not targeting exclusively mPFC-projecting vCA1 engrams, we could detect mCherry^+^ fibers in the mPFC (Fig. 3D), which is in line with the enrichment of vCA1 engrams in the projection to the mPFC (Fig. 1L). When assessing activity of GFP-labelled mPFC fear “A” engrams upon vCA1 engram “C” inhibition, we observed a reduction in their reactivation compared to VEH-injected controls (Fig. 3E), consistent with the notion of preferential connectivity between engram cells ^29^. Altogether, our results indicate that contextual hippocampal input controls mPFC engram reactivation during remote memory generalization.

### Activity of mPFC PV cells mediates fear engram recruitment during memory generalization

vCA1 neurons are known to directly contact PV interneurons in the mPFC and modulate different behaviours via feed-forward inhibition ^20,30^. In addition, recent work has revealed a role of PV interneurons in the reorganization of remote mPFC engrams during systems consolidation ^25^. Thus, we asked whether PV cells in the mPFC were differentially modulated by vCA1 inputs at remote recall. Using identical experimental settings as before, we observed contacts between mCherry^+^ fibers from vCA1 neurons and PV cells in the mPFC (Fig. S3C), consistent with previous work ^20,30^. Further, cellular quantifications revealed a higher proportion of active PV interneurons (PV^+^cFos^+^ / PV^+^) in AC when compared to AA animals for both TRAP2-Ai14 and VEH-injected *cFos::tTA* mice (Fig. 4A and S3D), and this parameter significantly correlated with engram recruitment into the recall ensemble across experiments (Fig. 4B). This included animals in which the vCA1–mPFC projection was inhibited via CNO-administration and which showed a concomitant reduction in PV activity (Fig. 4B and S3E), indicating a direct modulation of PV cells by vCA1 inputs.

**Figure 4:**
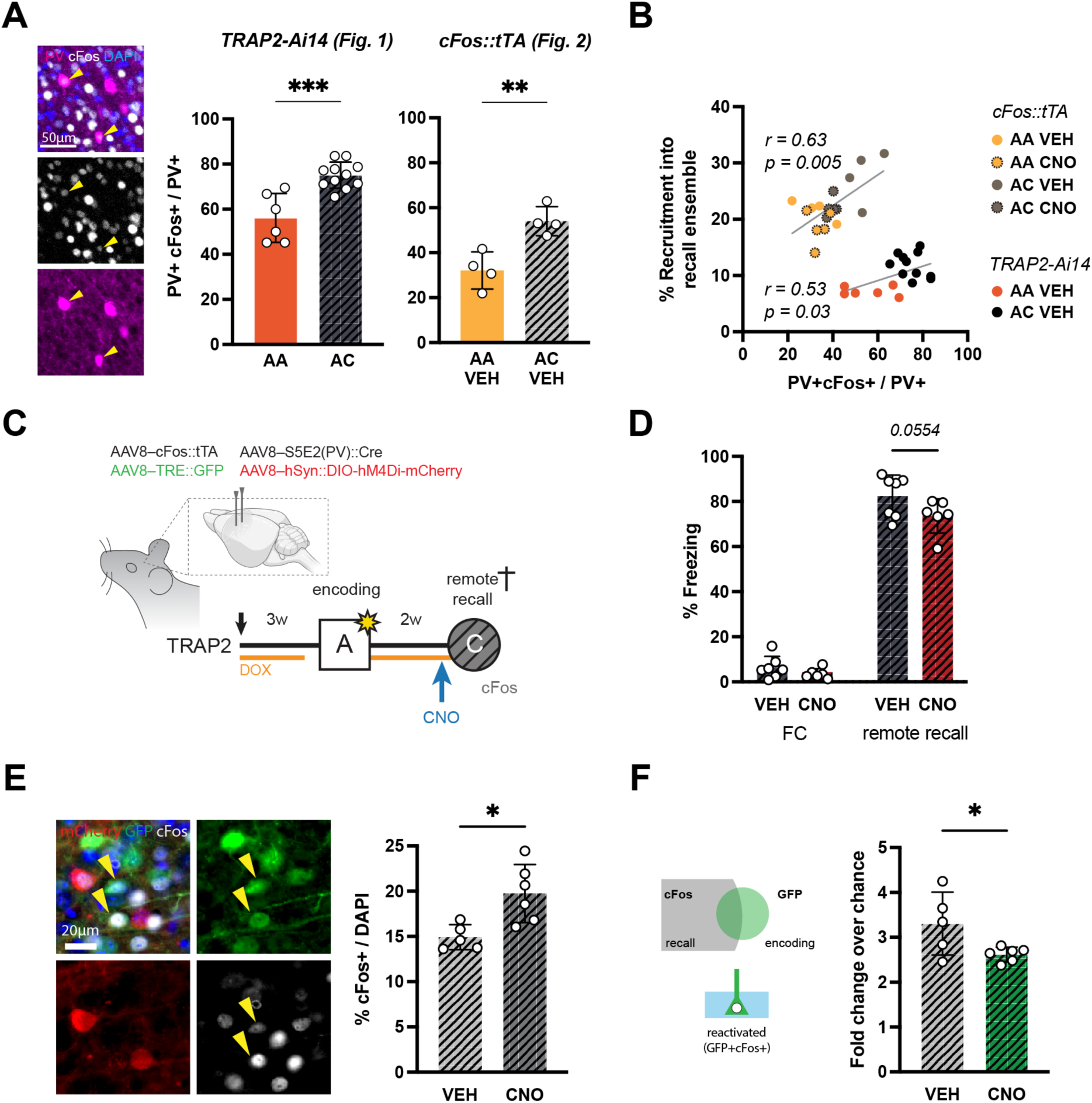
PV cells mediate mPFC fear engram recruitment during memory generalization. **A – B.** Immunofluorescence pictures and quantifications (A) of active mPFC PV cells in AA and AC mice (corresponding to the same animals as in Figs. 1 and 2, respectively). Note the increase in PV activity in AC mice in both transgenic lines, which significantly correlates with recruitment into the recall ensemble across experiments (TRAP2-Ai14 and *cFos::tTA*) and manipulations (VEH and CNO; B). **C.** Scheme depicting the experimental design for the manipulation of PV cells in the mPFC during memory generalization. In this setup, the indicated combination of viruses allows the expression of hM4Di-mCherry in mPFC PV cells, their inhibition via CNO administration and the additional tagging of fear “A” engrams. **D.** Percentage of freezing in AC mice injected with VEH or CNO, showing a reduction in memory generalization after inhibition of PV cells. **E – F.** Immunofluorescence picture with magnifications (E, left) of “A” (GFP^+^), recall (cFos^+^) and reactivated (GFP^+^cFos^+^, yellow arrowheads) mPFC ensembles, together with manipulated PV cells (mCherry^+^) and quantifications showing an overall increase in cortical activity (cFos^+^, E, right) and, conversely, reduced engram reactivation (F) in CNO as compared to VEH mice. Data are represented as bar graphs (filled or stripped for AA and AC mice, respectively) with individual animal values (dots; N > 5 for fear memory, > 3 for cellular analysis) and as correlation (B). Error bars indicate SD. Scale bars = 50 and 20 μm as indicated. *** *p* < 0.001; ** *p* < 0.01; * *p* < 0.05; ns, non-significant, by tests as indicated in Table S1. See also Figure S3

To causally evaluate whether PV cells mediate the effects of vCA1 inputs on fear engram recruitment, we chemogenetically inhibited PV cells in the mPFC. Here, the expression of Cre was coupled to an enhancer specific of PV interneurons ^31^ (Fig. S3F), which then became inhibited upon CNO administration 30 min before remote exposure to context C (Fig. 4C). This resulted in decreased freezing (Fig. 4D), confirming a role of PV interneurons in remote memory generalization. Further, we found that while inhibition of PV cells increased overall cortical activity (Fig. 4E), it conversely reduced reactivation of the fear “A” engram (Fig. 4F), suggesting a specific role of PV cells in regulating engram reactivation. Altogether, these results indicate that selective control of engram recruitment by vCA1 inputs occurs via modulation of PV cells during remote memory generalization.

## DISCUSSION

Here, we show that contextual vCA1 to mPFC inputs are necessary for time-dependent memory generalization. vCA1 has been proposed to dynamically encode a broader range of stimuli relevant to the animal than its dorsal counterpart ^32^, including contextual information ^20,33–37^. The routing of such stimuli to the mPFC, which acts both as a depository for long-term memories ^4,22^ as well as a key hub for decision-making and behavioural flexibility ^38,39.38,39^, could help the cortex to resolve whether generalization should or not occur. On this basis, we hypothesize that vCA1 acts as a contextual comparator that informs the mPFC about the presence in an environment similar enough to the fear-associated context and, consequently, triggers generalization for adaptive reasons. This would justify the presence of sustained reactivation in vCA1 engrams at remote times, since a representation of the original fear engram must be remembered and compared to the representation of the novel context. Thus, upon inhibition of the entire vCA1 region ^8^ or, more specifically, its mPFC-projecting cells or representation of context C as shown here, value information cannot be compared nor routed to the mPFC, explaining the decrease in generalization. This was not the case after inhibiting the vCA1 homecage ensemble, probably due to the bigger resemblance between A and C contexts, thus showing that the effect is context-specific.

Our data additionally demonstrate that such hippocampal routing of contextual information controls mPFC engram reallocation during generalization, and that this process is modulated by PV interneurons. Previous work described a reorganization of cortical circuits during consolidation that favoured the recruitment of cells not originally active during learning when animals were tested in the fear-associated context at remote times ^23,24^. In contrast, our results show an increased relative contribution of fear engrams to the recall ensemble during memory generalization in a novel context, likely contributing to a heightened freezing response. This effect was prevented when vCA1 inputs or mPFC PV cells were acutely inhibited before recall, suggesting that generalization emerges as a flexible reconfiguration of the consolidated hippocampal-cortical circuit. Supporting this notion, approaches destabilizing or stabilizing hippocampal circuits have been shown to increase or decrease remote memory generalization, respectively ^26,40^, presumably by locking networks into more or less flexible states. Our results also underscore PV cells as key mediators of such circuit remodelling, not only at early stages of consolidation as recently shown ^25^ but also acutely at remote times. Remarkably, this flexible modulation of activity is engram-specific, as evidenced in our PV manipulation experiments resulting in opposite effects in the activity of cortical engram and non-engram cells. Such specificity evokes previous work on inhibitory control of DG ensembles during recent memory recall ^41–43^, but how interneurons compute different contextual inputs and distribute activity in memory-relevant ensembles remains to be explored.

Finally, our results can also be interpreted from a systems perspective and in the light of the different theories for remote memory. Our findings are reminiscent of the concept of memory indexing in that hippocampal ensembles control the recruitment of cortical engrams to drive remote memory expression. Interestingly, it is not the original hippocampal ensemble that was needed for remote recall, indicating that the mPFC no longer needs the hippocampal index for recall of the fear-associated context. Instead, it was the contextual information of the novel context that promoted the recruitment of the past fear cortical engram, triggering memory generalization. In this context, our work provides an experimental example of how “fluid” ensembles ^44^ or memory transience ^1^ sustain adaptability and flexible behaviours. This expands our understanding of the role of the hippocampus in remote memory and supports the notion that time-dependent generalization is an active, rather than a passive loss-of-precision, process ^7^.

### Limitations of the study

Given the complex projection patterns of vCA1 neurons ^10^, whether and how the manipulation of mPFC-projecting cells affects other brain areas through axon collaterals, as well as other kinds of memory such as social or spatial, was not studied. In addition, our inhibition manipulations relied on chemogenetic as opposed to optogenetic approaches, thus lacking fine temporal resolution that would be needed to ascertain the influence of vCA1 inputs in early vs. late phases of generalization. Finally, dilution of GFP expression for engram identification beyond the 2 weeks timepoint prevented us from studying generalization at even later stages.

## Supporting information

Table S1: Statistics

## Acknowledgements

The authors would like to thank Lisa Watt, Austeja Dapkute and Alice Valérie Juanico for technical assistance, Bernard Schneider for help with viral vectors, the EPFL’s Center of Phenogenomics for ensuring animal welfare, the Histology Core Facility for support with brain sectioning and staining and the Bioimaging and Optic Core Facility for their expertise during image analysis. The authors would also like to thank all present and past members of the Gräff lab for scientific feedback.

## Author contributions

This work was conceptualized by GBV and planned together with JG. GBV performed and analysed all experiments supported by IF, JO and MS. The paper was written by GBV and JG and commented by all authors. Correspondence and requests for materials should be addressed to JG.

## Data and materials availability

All data are available in the main text and supplemental information.

## METHODS

### EXPERIMENTAL MODEL

All animals and procedures were approved by the Veterinary Office of the Federal Council of Switzerland under the animal experimentation licenses VD3662.

#### Animals and procedures

TRAP2 (*Fos::Cre^ERt2^*), TRAP2-Ai14 (*Fos::Cre^ERt2^ x ROSA-tdTomato*) and *cFos::tTA* mouse lines were bred in house in the C57Bl/6JR background, and 2-4 months old transgenic offspring of either sex was used as experimental mice. Animals were group-housed in a 12h light/dark cycle with water and food available *ad libitum,* habituated to the experimenter for 2-3 days prior to the experiments and randomly assigned to experimental groups. For engram labelling, 4-hydroxytamoxifen (tam) was prepared as previously described ^23^ and injected in TRAP2 mice intraperitoneally (50mg/kg) immediately after conditioning. Alternatively, doxycycline (Dox) was provided through the drinking water (0.2mg/ml) at least 7 days before surgeries and only ceased 2 days before the opening of the labelling window for engram tagging, after which it was immediately reintroduced. For chemogenetic inhibition, CNO was prepared as previously described ^45^ and injected intraperitoneally (3mg/kg) 30 minutes before remote recall.

### METHODS DETAILS

#### Viral constructs and stereotaxic injections

cFos::tTA (gift from William Wisden, addgene plasmid #66794 ^46^) viruses were generated in house (EPFL’s Bertarelli Foundation Gene Therapy Platform) in the AAV8 serotype at a titer of 2.8×10^13^ VG/ml. AAV8–TRE::GFP viruses were provided by UNC Vector Core, at a titer of 4.1×10^12^ VG/ml. AAVretro–Pgk::Cre viruses (1.7×10^13^ VG/ml) were a gift from Patrick Aebischer (Addgene viral prep #24593-AAVrg ; RRID:Addgene_24593). AAVretro–CAG::GFP (7×10^12^ VG/ml) viruses were a gift from Edward Boyden (Addgene viral prep #37825-AAVrg; RRID:Addgene_37825). AAV8–hSyn::DIO-hm4Di-mCherry (1.8×10^13^ VG/ml) was a gift from Bryan Roth (Addgene viral prep #44362-AAV8; RRID:Addgene_44362). AAV1–CAG::FLEX-EGFP was a gift from Hongkui Zeng (Addgene viral prep #51502-AAV1; RRID:Addgene_51502). The PV-specific enhancer S5E2 (a gift from Jordane Dimidschstein; Addgene plasmid #135631; RRID:Addgene_135631) was subcloned into a Cre-containing backbone and viruses were generated in house (EPFL’s Bertarelli Foundation Gene Therapy Platform) in the AAV8 serotype at a titer of 1.2×10^13^ VG/ml.

For stereotaxic surgeries, mice were anaesthetised with a mix of fentanyl (0.05 mg/kg) midazolam (5 mg/kg), and metedomidin (0.5 mg/kg) injected intraperitoneally and followed by subcutaneous injection above the head of an analgesic mix including lidocaine (6 mg/kg) and bupivacaine (2.5 mg/kg). Mice were placed in the stereotaxic frame (Kopf Instruments), their skin disinfected and opened. Holes were drilled on the adjusted skulled according to the injection area. Viruses were injected bilaterally through pulled micropipettes (Blaubrand, intraMARK, 708707) at a rate of 0.2 μl/min in the following coordinates and volumes: vCA1, AP -3.16mm, ML ± 3.25mm, DV -4.25, 4 and 3.75mm, with 200, 50 and 50nl per injection site; mPFC, AP +2.0mm, ML ± 0.35mm, DV 2.25mm; 250nl per side. Needles were left at the injection site for 5 minutes and slowly pulled up to prevent backflow. Wounds were sutured and mice given atipamezole (2.5mg/kg) to reverse anaesthesia, after which they were placed in a heated cage with painkiller in the drinking water (Dafalgan 2mg/ml) and wet food in the bedding for recovery. Behavioural experiments started no sooner than 2 weeks after surgeries. Mice with poor transduction efficiency (< 2% positive cells) or clearly mistargeted were excluded from further analyses.

#### Fear conditioning

Fear conditioning was performed using a Multi Conditioning System (TSE)^45^. On the day of conditioning, mice were acclimatized to the behavioural room for 30min and then allowed to explore the conditioning box for 3 min in a first habituation phase. After this, mice received electric foot-shocks (3x0.8mA, 2s long and separated by 8s intervals) and were moved to a holding cage 15s later. Recall occurred 2 days or 2 weeks afterwards (recent or remote, respectively), by exposing mice to the same conditioned (A) or a novel (C) context for 3 min without shocks. Context A consisted of a squared shuttle box with transparent walls and metal grid, ventilated, non-illuminated and cleaned with 5% ethanol. Context C consisted of a round shuttle box with black-white striped plexiglass walls and plastic floor, lit from above, no ventilation and cleaned with 5% ethanol lightly scented with Virkon. Freezing was used as a proxy for fear memory and quantified automatically by infrared beam cutting detection using 1s of immobility as threshold.

#### Open field

Effects in anxiety and exploratory activity were measured using an open field test. Briefly, mice were moved to the testing room for 2 days before behaviour for habituation. On the test day, mice were allowed to freely explore a circular arena (20 cm radius) lightly lit from above for 10 minutes, after which they were moved to a holding cage. Sessions were recorded from above and tracked using Ethovision (Noldus). Time spent at the center vs. periphery was calculated throughout the test and used as a measure of anxiety. The arena was cleaned between trials with 5% Mucasol.

#### Histology

Mice were administered a lethal dose of pentobarbital (150mg/kg) 90 minutes after the last recall session and transcardially perfused with PBS followed by 4% paraformaldehyde (PFA) in PBS. Extracted brains were further fixed by overnight immersion in PFA and cryoprotected 3 days later in 30% sucrose in PBS. Sections were cut at 30 µm in a cryostat and serially collected and preserved in antifreezing solution (30% ethylene glycol, 15% sucrose, 0.02% azide in PBS) until staining. For stereological quantifications, one every six sections per brain area (for total of 6-10 sections) were used for immunofluorescence. Sections were first blocked and permeabilized in blocking solution (1% BSA, 0.3% triton in PBS) for 2h and then incubated with primary antibodies overnight at 4°C. The primary antibodies included rabbit anti-RFP (Rockland 600-401-379, 1:2000), goat anti-GFP (AB6673, 1:1000), rabbit anti-GFP (AB290, 1:1000), mouse anti-GFP (AB1218, 1:1000), guinea-pig anti-cFos (Sysy 226308, 1:4000), mouse anti-PV (Swant Cat# 235, 1:5000). The following day, sections were washed three times in PBS and then incubated with secondary antibody (Invitrogen or Jackson Immunoresearch, 1:800) in blocking solution for 3h. Finally, sections were washed in PBS, counterstained with DAPI (1:5000 in PBS, Invitrogen H3570) and washed three more times before mounting on glass slides and covered with Fluoromount-G medium to preserve fluorescence.

### QUANTIFICATION AND STATISTICAL ANALYSIS

#### Imaging and quantifications

For cellular quantifications, images from at least 3 sections per animal were acquired using an Olympus SlideScanner VS120 L100 with 10x (for mPFC, 0.65 µm pixel size) or 20x (for vCA1, 0.32 µm pixel size) objectives. Quantifications in vCA1 were performed from Bregma –2.9 to –3.8 mm, whereas mPFC quantifications were restricted to the anterior cingulate, prelimbic and infralimbic regions of the mPFC, rostral to the callosal junction, from Bregma 2.25 to 1.45mm. Cell numbers for engram reactivation analyses were extracted using QuPath ^47^ automatic cell detection and co-localizations scripts. Cells positive for individual channels (e.g., Tom^+^, GFP^+^, cFos^+^) were first detected following manually pre-set thresholds relative to the background noise and later classified as double or triple positive by proximity. Total numbers of cells were estimated via detection of DAPI cells, which was then used to calculate densities and chance probabilities. Cells were normalized either by the total number of a given subpopulation (e.g., Tom^+^ or GFP^+^, for reactivation of encoding engram) or by chance (total / chance; total = double positive / DAPI; chance = [single positive *a* / DAPI] * [single positive *b* / DAPI]).

#### Statistics and reproducibility

Quantification of cell numbers were performed on independent biological replicates (N = mice) and depicted as means ± SDs, correlations or line-plots as indicated in the figure legends. Statistical analyses were performed in Prism 10 (GraphPad) and significance was assessed by two-tailed, unpaired (different animals) or paired (same animal or group) Student’s t-test and ANOVA (one or 2-way) or non-parametric version. Behavioural analyses were performed on 5–17 mice per group (as indicated in figure legends) from at least 2 different batches, and data is depicted as bar graphs. Statistical significance was calculated by repeated measures two-way ANOVA.

---

**Figure S1:**
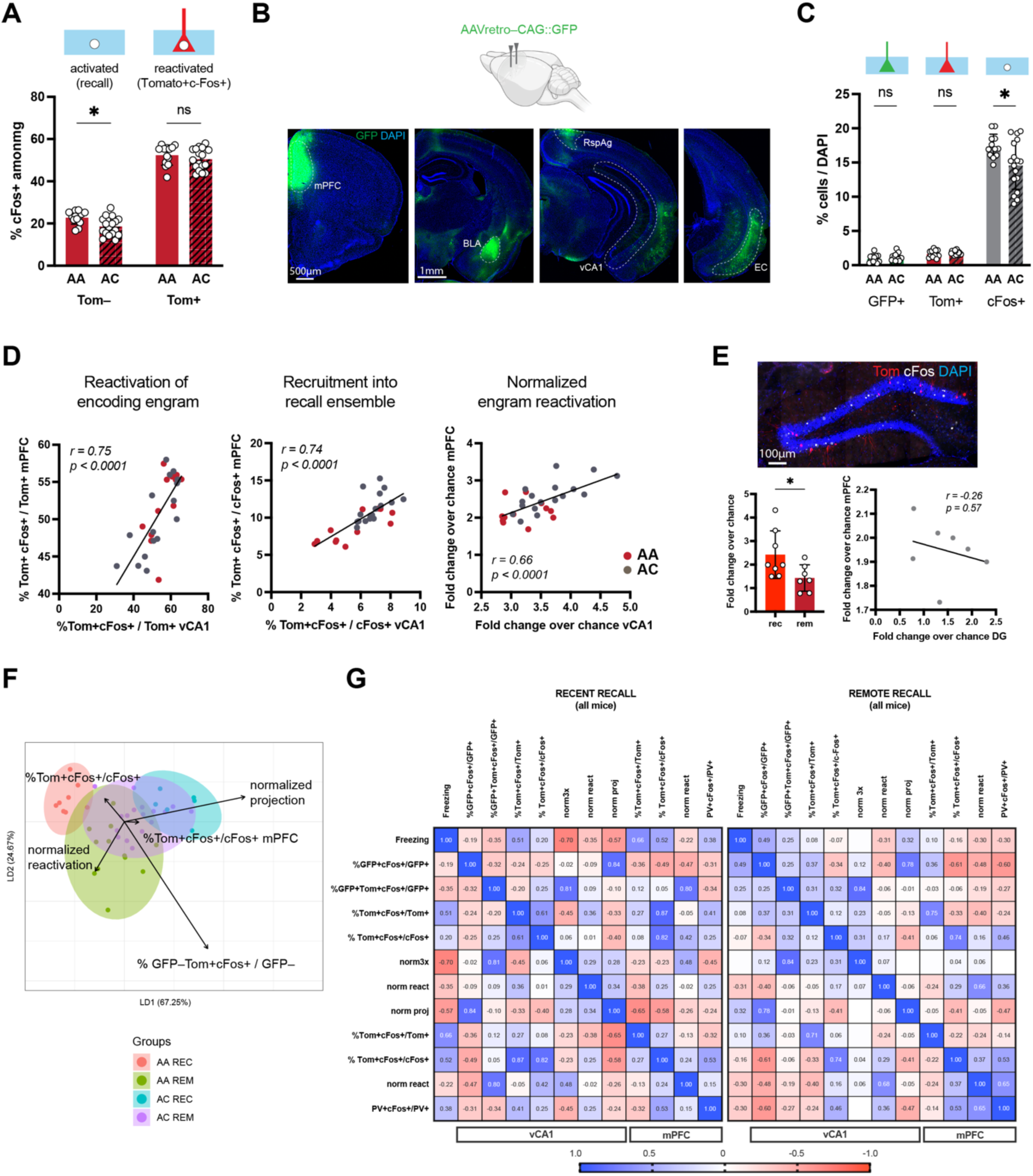
Cellular changes associated with remote memory generalization, related to Figure 1. **A.** Percentage of active mPFC cells among non-engrams (Tom^−^) and engrams (Tom^+^) of AA and AC TRAP2-Ai14 mice. **B.** Immunofluorescence pictures showing canonical mPFC inputs upon injection of a retrograde tracer (GFP^+^), including the basolateral amygdala (BLA), vCA1, retrosplenial agranular (RspAg) and entorhinal cortex (EC). **C.** Cell densities in vCA1, including percentage of traced (GFP^+^), engram (Tom^+^) and recall (cFos^+^) cells of AA and AC mice. **D.** Correlation between mPFC and vCA1 engram reactivation parameters of AA (red) and AC (grey) mice. **E.** Immunofluorescence picture (top), and quantification (total over chance, bottom left) of engram reactivation in the DG of AA mice tested at recent (light red) and remote (dark red) times, showing time-dependent decrease in reactivation and lack of correlation with mPFC activity at remote times (bottom, right). **F.** Linear discriminant (LD) analysis of all cellular parameters quantified in AA and AC mice (individual dots), tested at recent or remote times (colour coded as indicated). Note the difference in cluster overlap between recent and remote groups, as well as the greater contribution of engram (e.g., “%Tom^+^cFos^+^/cFos^+^”) or projection (e.g., normalized projection) parameters to the separation of AA or AC points, respectively. **G.** Heatmap showing the correlation between freezing, vCA1 and mPFC cellular parameters of AA and AC animals tested at recent and remote times. Cellular parameters include percentages over labelled projections (GFP^+^), fear engram (Tom^+^), recall (cFos^+^) and PV cells (PV^+^), as well as normalized by chance levels (for projection -norm proj-, reactivation -norm react- and reactivated projections -norm3x-). Positive and negative pearson correlation values are represented in blue and red squares, respectively. Note the positive correlation between freezing and engram parameters at recent, as well as with projection parameters at remote, times. Data are represented as bar graphs (filled or stripped for AA and AC mice, respectively) with individual animal values (dots; N > 6) and as heatmaps with pearson correlation value (G). Error bars indicate SD. Scale bars = 1mm, 500 μm and 100 μm as indicated. * *p* < 0.05; ns, non-significant, by tests as indicated in Table S1.

**Figure S2:**
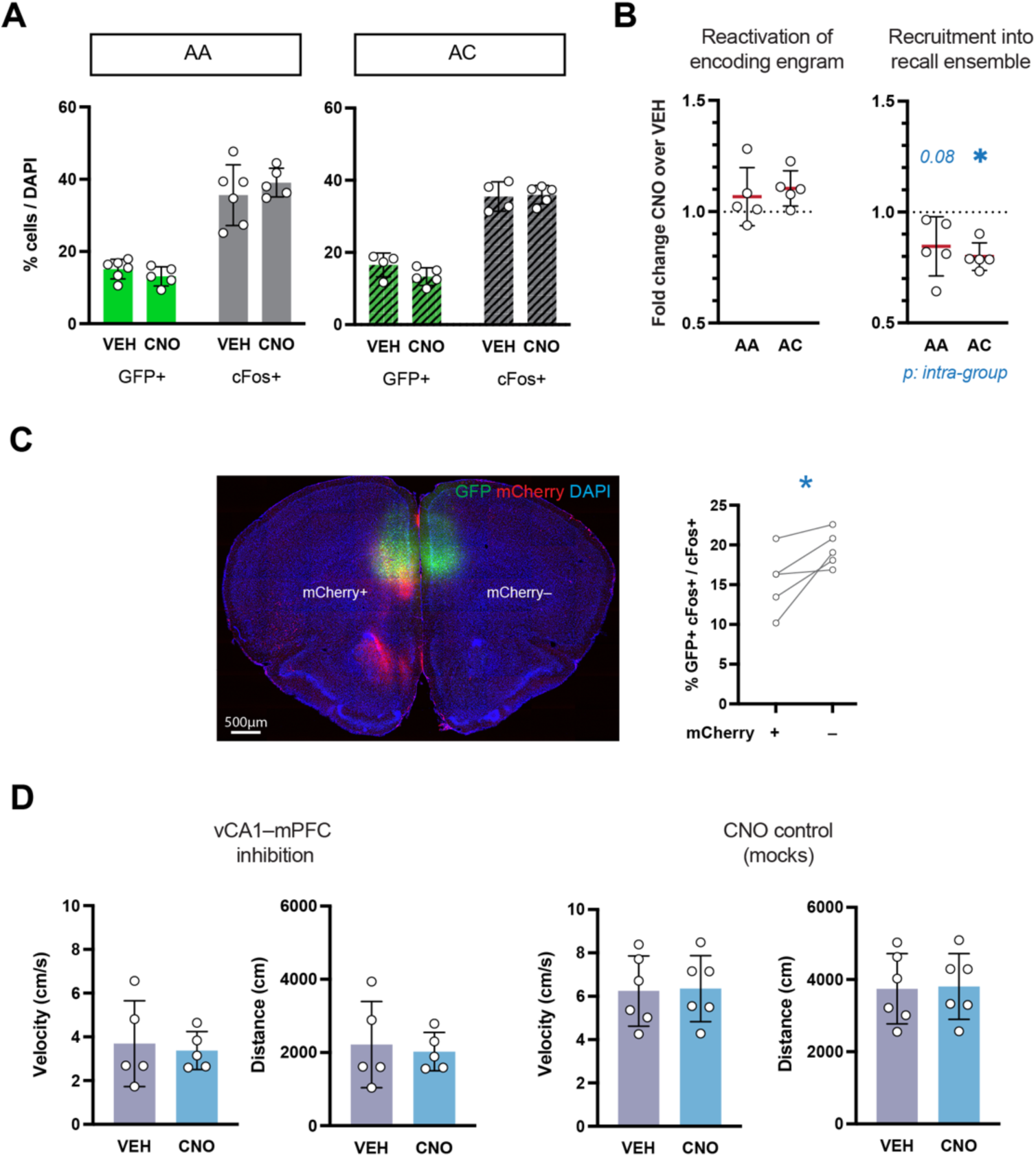
Control for vCA1–mPFC projection inhibition, related to Figure 2. **A.** Cell densities in the mPFC of VEH and CNO-injected AA (filled) and AC (stripped) mice. **B.** Quantification of reactivation of encoding engram (left) and recruitment into the recall ensemble (right) of AA and AC mice, as indicated. **C.** Immunofluorescence (left) and quantification (right) showing contralateral differences in fear engram recruitment between inhibited (presence of mCherry^+^ fibers) vs. non-inhibited (mCherry^−^) sides. **D.** Quantification of exploratory parameters (velocity and distance travelled, as indicated) between VEH (grey) and CNO (blue) animals with inhibited vCA1–mPFC projection (left) and, as control for CNO administration, non-injected *cFos::tTA* mice (right). Data are represented as bar graphs with individual animal values (dots; N > 3 for cellular analysis, A; > 4 for behaviour, D), as fold change of the CNO animals over the average of their respective VEH-injected group, with blue significance indicating differences between VEH and CNO mice (B), and as intra-animal reactivation comparisons (C). Error bars indicate SD. Scale bars = 500 μm. * *p* < 0.05; ns, non-significant, by tests as indicated in Table S1.

**Figure S3:**
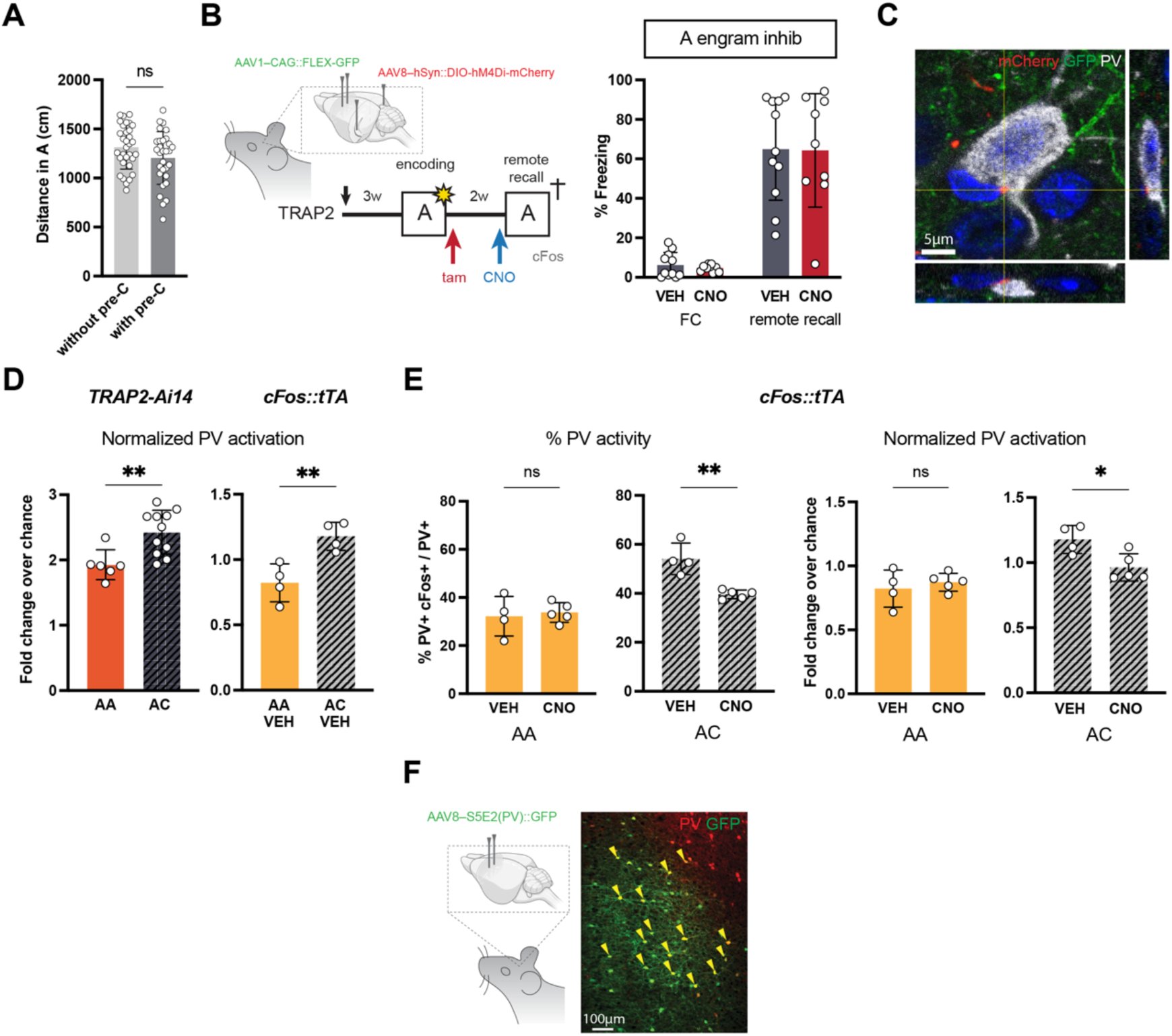
Control for vCA1 engram and PV inhibition, related to Figures 3 and 4. **A.** Distance travelled during the habituation phase in the context A of animals tested without (light grey) and with (dark grey) a previous exposure to context C. **B.** Scheme depicting the experimental design for the manipulation of vCA1 fear ensembles during memory generalization (left) and freezing quantification (right), indicating that vCA1 “A” engrams are not needed for remote recall in the fear-associated context. **C.** Confocal magnification showing physical contacts between vCA1 fibers (mCherry, in red) and mPFC PV (in white), with orthogonal projections. **D – E.** Quantification of PV activity in TRAP2-Ai14 (as in Figure 1) and *cFos::tTA* (as in Figure 2, VEH and CNO) mice, exposed to the same conditioned (AA) or novel (AC) context, and showing hippocampal-dependent increased PV activity in AC mice. Data is shown as percentage over PV population or normalized by chance, as indicated. **F.** Scheme depicting the injection of S5E2(PV)::GFP (left) and immunofluorescence (right), showing expression of GFP specifically in PV cells in the mPFC. Data are represented as bar graphs (filled or stripped for AA and AC mice, respectively) with individual animal values (dots; N > 8 for behaviour, > 3 for cellular quantifications). Error bars indicate SD. Scale bars = 100 and 5 μm as indicated. ** *p* < 0.01; * *p* < 0.05; ns, non-significant, by tests as indicated in Table S1.

